# Dual orexin antagonist DORA-22 suppressed posttraumatic seizures and enhances GABAergic inhibition in dentate granule cells

**DOI:** 10.1101/2022.04.17.488582

**Authors:** Sruthi R. Konduru, Jesse R. Isaacson, Zihao Zhou, Rohan K. Rao, Danny J. Lasky, Swati S. Vattem, Sophie J. Rewey, Mathew V. Jones, Rama K. Maganti

## Abstract

**Background:** Traumatic brain injury (TBI) can result in posttraumatic epilepsy (PTE) and sleep disturbances. We hypothesized that treatment with sleep aids after TBI can ameliorate PTE.

**Methods:** CD-1 mice underwent controlled cortical impact (CCI), sham craniotomy or no craniotomy. Sham and CCI groups underwent a month-long treatment with sleep aids including a dual orexin antagonist (DORA-22) or THIP (gaboxadol). We performed week-long EEG recordings during week-1 of treatment and again at months 1, 2 and 3. Seizure analysis occurred at all-time points and sleep analysis occurred in week-1 and month-1 recordings in all groups. Subsets of animals in sleep aid-treated and untreated CCI, and sham groups were subjected to voltage clamp experiments.

**Results:** DORA-22 treated group had seizures at week-1 but none at months 1-3. TBI reduced amplitude and frequency of miniature inhibitory synaptic currents (mIPSCs) in dentate granule cells and these changes were rescued by DORA-22 treatment. Sleep analysis showed that DORA-22 increased non-rapid eye movement sleep (NREM) in the first 4hours of lights-off whereas THIP increased REM sleep in the first 4-hours of lights-on in week-1, At month-1 both treatments reduced time in NREM during lights-off. TBI increased NREM delta power (NΔ) along with loss of the homeostatic overnight decline of NΔ in week-1 regardless of treatment. DORA-22 and THIP treatment restored NΔ to levels similar to no craniotomy animals at month-1.

**Conclusions:** DORA-22 treatment suppressed posttraumatic seizures possibly due to enhanced GABAergic inhibition in dentate granule cells. DORA-22 may have therapeutic potential in suppressing PTE.

**Summary for Social Media:** Traumatic brain injury (TBI) can result is posttraumatic epilepsy and sleep disturbances. There are no treatments to prevent these complications. We tested whether treatment with sleep aids after TBI can mitigate seizures and sleep disturbances. We found that a sleep aid DORA-22 but not THIP suppressed post traumatic seizures possibly be enhancing GABAergic inhibition in the hippocampus. Sleep analysis showed that TBI disrupts the sleep homeostatic drive and DORA treatment restored it. Findings may lead to potential disease modifying therapy for posttraumatic epilepsy.

## Introduction

An estimated 2.87 million people experience TBI, with over 280,000 hospitalization each year in the United States^1^. Sequelae of TBI include posttraumatic epilepsy (PTE), sleep disorders, cognitive and motor deficits, and mood disorders such as depression or posttraumatic stress disorder (PTSD)^2^. Epidemiological data show that PTE constitutes 5% of all cases and over 20% of cases of acquired epilepsy^4^. Posttraumatic seizures can occur early or late and 80% of posttraumatic seizures begin in the first 2 years after TBI but latency can be years or even decades^5^. Epidemiological evidence also shows that sleep disturbances can begin immediately after TBI and persist long after the injury^6^. Sleep complaints in TBI patients range from insomnia to hypersomnia^7^ with polysomnographic studies showing sleep fragmentation and multiple sleep latency tests showing reduced latencies^8^.

Sleep disturbances in TBI result from injury to regions such as thalamus, hypothalamus and brain stem^9^ and PTE from injury to hippocampus and cortex^10^. In mouse models of TBI, PTE occurs in 25-33% of mice^10-11^ which is similar to our experience^12^. Data on sleep disturbances in mouse models of TBI showed hypersomnia, increased NREM sleep and short wake bouts acutely or chronically^13-14^. In a controlled cortical impact (CCI) model of TBI, we observed no changes in sleep architecture acutely or in the first month after TBI but increases in NΔ was seen acutely that persisted 1-month post-TBI^12^.

Orexins are hypothalamic neuropeptides that regulate arousal and wakefulness. Physiologic functions of orexins are mediated by two G-protein coupled receptors namely type 1 (Orx1) and type 2 (Orx2)^15^. Orexin receptor antagonists are known to improve sleep and are approved for treatment of insomnia^16^. Orexins are also known to be involved in synaptic plasticity and long-term potentiation in the hippocampus^17^ mediated via glutamatergic as well as GABAergic receptors^18^. There are several reports which demonstrated that orexin antagonists suppress seizures in mouse models of epilepsy^19-21^, but the exact mechanisms are not understood.

Sleep disturbances and PTE often occur in the same patient after TBI. Moreover, sleep disruptions and epilepsy have a complex bidirectional relationship where each can exacerbate the other^22^. We hypothesized that enhancing sleep immediately after TBI may be protective against development of PTE and tested it by administering 2 different sleep aids: a dual orexin antagonist (DORA-22) and an agonist of δ subunit-containing GABA_A_ receptors (THIP), daily for 1-month after CCI. We found that the former but not the latter, suppressed PTE. Moreover, TBI diminished GABAergic inhibition in dentate granule cells and was rescued by month-long DORA-22 treatment. Sleep NREM delta power, a marker of sleep homeostatic drive which increases in proportion to preceding wakefulness but rapidly declines with sleep^23^, was increased by TBI and is restored by a month-long treatment with both DORA-22 and THIP. Moreover, TBI altered the homeostatic overnight decline in NΔ regardless of treatment at week-1. THIP treatment did not restore this homeostatic overnight decline in NΔ at month-1.

## Methods

### Animals

Adult male CD-1 mice (∼4 months old) were obtained from the Jackson Laboratory (Bar Harbor, ME). Mice were singly housed in their recording chambers during electroencephalography (EEG) recording days and were group-housed otherwise. Mice were housed under a 12:12-h light:dark cycle with access to food and water ad libitum. All procedures involving the use of animals were approved by the University of Wisconsin IACUC in accordance with the US Department of Agriculture Animal Welfare Act and the National Institutes of Health Policy on Humane Care and Use of Laboratory Animals.

### TBI Induction and Electrode Implantation

CCI was performed according to methods described previously^12^. Briefly, under isoflurane anesthesia, a 4-5 mm craniotomy was performed in a stereotaxic frame and moderate-to-severe CCI was performed using a Precision Cortical Impactor (Hatteras Instruments, NC) (2 mm depth, 5 m/s impact velocity, and 100-200 msec dwell time) (n=40). The sham injury group (n= 16) had a craniotomy with no CCI while a control group had no craniotomy or CCI (n=6). All groups received right frontal and left parietal epidural screw EEG electrodes (touching the dura) and 2 stainless steel wires implanted in nuchal muscles for EMG, at the same time as CCI.

### Analysis of Seizures, Vigilance States and NREM Delta Power

Video-EEG data was manually reviewed in 1-second epochs to identify seizures. After obtaining absolute seizure counts, a normalization was used per day of recording per animal. EEG data were manually scored for vigilance states, in 4-second epochs using Sirenia Sleep (Pinnacle Technologies, Lawrence, KS), divided into wake, NREM, or REM using EEG and EMG trace waveforms according to methods described previously^12^. All epochs with seizures and artifactual EEG were excluded from sleep analysis. Day 4 or Day 5 of recording was selected randomly for sleep scoring. Power in different frequency bands was calculated for each scored 4-second NREM epoch of frontal EEG by establishing power spectral density with fast Fourier transformation and integrating the power spectral density of the delta band (0.5-4 Hz). To account for inter animal variability, delta power values were normalized to the sum of power integrated over delta (0.5-4), theta (5-7 Hz), alpha (8-13), beta (14-23 Hz) and gamma (30-70 Hz) bands. Custom MATLAB scripts were used to calculate the normalized NΔ between different treatment groups.

### Drugs treatments

Sham injury and CCI groups underwent drug treatment with a DORA called DORA-22 (a gift from Merck, Inc) at 100 mg/kg with 20% TPGS (tocopherol with polyethylene glycol) as vehicle (CCI; n=12; sham; n=8) and THIP (4,5,6,7-tetrahydroisoxazolo[5,4-c]pyridin-3-ol) at 2 mg/kg with PBS (phosphate-buffered saline) as vehicle (CCI n=12; sham n=8). Drug treatments were performed daily via oral gavage approximately at ∼8-9 am each morning (shortly after lights-on or the time of sleep onset for rodents).

### Patch-Clamp Electrophysiology

Patch-clamp recordings and analyses were performed according to methods described previously^24^. Briefly, subsets of sham, CCI vehicle treated and DORA treated animals were sacrificed at the end of month-long EEG recording or drug treatment to obtain horizontal hippocampal slices. The procedure described previously was followed and miniature inhibitory postsynaptic currents (mIPSCs) were recorded under voltage-clamp of -70Mv and detected using the MiniAnalysis program (Synaptosoft, Decatur, GA)^24^.

### Data Analysis

Differences in seizure frequency normalized to number of days of recording and number of animals at each time point (week-1 and months 1-3) and the 24-hour mean NΔ between treatment groups was analyzed using one-way ANOVA and post hoc Tukey tests for multiple comparisons. Two-way ANOVA with repeated measures was used for analysis of vigilance states (2-hour blocks) and for NΔ (1-hour blocks) across the 24-hours to understand the interactions of time bin as an independent variable and treatment as an independent variable with appropriate post-hoc corrections. Differences in mean NΔ in the first 3 hours (initial portion of rodent sleep) and the last 2 hours (towards the end of the night) was also compared using paired t-tests. All statistical analyses were performed in Graph Pad Prism with an alpha value of 0.05. Electrophysiology data was analyzed with home written scripts in MATLAB and effects of treatment on mIPSC amplitude and frequency was assessed with Kruskal-Wallis test.

### Data Availability

The custom MATLAB scripts utilized for NREM delta power analysis as well as data generated will be available on request.

## Results

### 1. DORA treatment for 1-month post-CCI suppressed posttraumatic seizures

Seizures were seen in ∼20% of CCI group (16/64 animals) and not in sham group or the no craniotomy controls. Absolute seizure counts varied from 0-44 in a given week in the treatment groups. Analysis of normalized seizure counts showed that the DORA-22 treated group had seizures only at week 1 and not at months 1-3, whereas TBI alone and TBI plus vehicle, and THIP treated TBI groups had seizures at all the time points (Table 1). Seizure characteristics were similar to what we reported previously^13^.

**Table 1:** The normalized daily seizure frequency in groups of animals with CCI that had no drug or vehicle, DORA-22 and THIP treatments. 2-way ANOVA showed that there was not a statistically significant interaction of time point and seizure frequency but treatment group had a significant interaction (F (3, 6) = 1.718; p=0.01) with post hoc Tukey test showing significant difference between DORA and THIP treated groups.

|  | Week 1 | Month 1 | Month 2 | Month 3 |
| --- | --- | --- | --- | --- |
| TBI alone or<br>TBI+ Vehicle<br>(n) | 1.21 (n=41) | 1.98 (n=24) | 2.78 (n=24) | 1.88 (n=21) |
| TBI+DORA (n) | 2.78 (n=14) | 0 (n=10) | 0 (n=8) | 0 (n=8) |
| TBI+THIP (n) | 8.33 (n=12) | 4.17 (n=12) | 4.69 (n=12) | 3.19 (n=8) |

### 2. TBI impaired GABAergic inhibition in dentate granule cells and was rescued by DORA-22 treatment

Seizures are thought to result from an imbalance of excitation and inhibition^25-26^ and increasing inhibition is favorable for neuronal/network excitability. Proposed mechanism of PTE is loss of synaptic inhibition due to loss of hilar interneurons in dentate granule cells^27-28^. In addition, prior work showed that DORA-22 treatment lowered glutamatergic excitation and increased GABAergic inhibition^18^. Therefore we chose to examine GABAergic inhibition in dentate granule cells by measuring mIPSCs in TBI or sham groups. Following EEG recordings at the end of 1-month, subsets of vehicle treated and DORA-22 TBI groups as well as sham injury group were sacrificed and horizontal hippocampal slices were obtained for voltage-clamp recordings in dentate granule cells (Figure 1), to evaluate the effects of TBI on synaptic inhibition (8 slices from 5 animals in each group). We found that the amplitude and frequency of mIPSCs were reduced in TBI compared to Sham or vehicle treated TBI. DORA-22 increased mIPSC amplitude and frequency in TBI group (Figure 3). Thus some of the consequences of TBI may be mediated by disrupted GABAergic inhibition and DORA-22 treatment rescued it.

**Figure 1:**
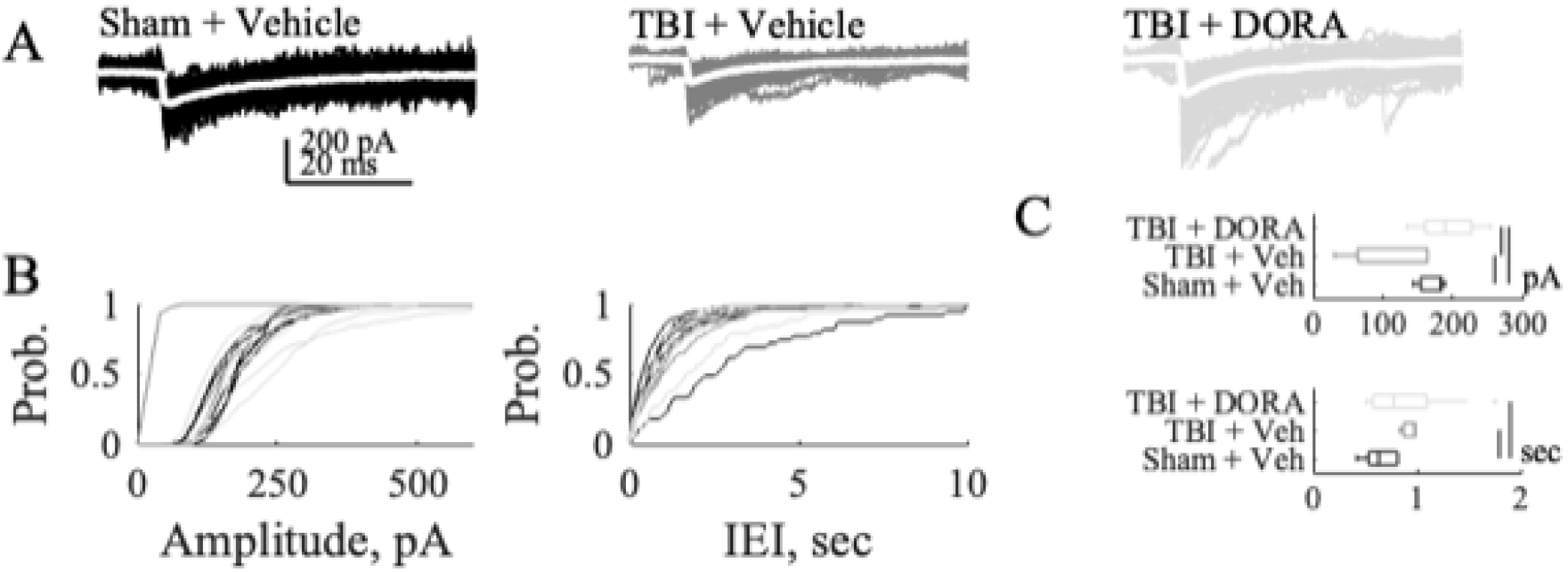
TBI effects on mIPSCs are partially rescued by a dual orexin antagonist (DORA-22): (A) Example mIPSCs from dentate granule cells. White overlays are the averages. (B) Cumulative amplitude and inter-event interval (IEI) distributions. Same color scheme as in A. (C) Box-and-whisker plots of median amplitudes (top) and IEIs (bottom). Vertical black bars indicate pairs of significant differences (p≤0.05, Kruskal-Wallis test).

### 3. DORA and THIP had different effects on sleep architecture

Given that DORA-22 and THIP are both sleep aids, we examined how TBI treatment with sleep aids altered sleep architecture. No differences were seen between DORA-22 treated (n=6), THIP (n=6) treated on TBI untreated or vehicle treated (n=12) in overall sleep efficiency (percent of time in sleep in 24 hours) [F (3, 7)=2.49; p=0.26], and sleep bouts (time spent each vigilance state) [F (3, 7)=2.25; p=0.24] or the total time spent in NREM [F (3, 6)=2.19; p=0.18] or REM [F (3, 6)=0.18; p=0.24] sleep across of the 24-hours at week 1 or month 1 and during lights-on (06:30-18:30 pm) or lights-off (18:30-06:30) periods (data not shown).

Analysis of vigilance states in 2-hour blocks across the 24-hours showed some changes in sleep architecture. At week-1, DORA treated group had greater NREM sleep in the first 4 hours of lights-off and REM sleep was greater in DORA-22 and THIP treated group in the first 4 hours of lights-on (Figure 2 A-B). At month-1 however, the time spent in NREM sleep was lower in the second 2-hour block of lights-on for THIP and higher for TBI only or TBI with vehicle during first two 2-hour blocks of lights-off. REM sleep was lower for both DORA-22 and THIP treated groups at month-1 in the second 2-hour block (Figure 2 C-D). Most importantly however, the diurnal oscillation of NREM and REM sleep (initial increase in lights off followed by a gradual decline and later a gradual increase) was better preserved in all groups at month 1 (Figure 2).

**Figure 2:**
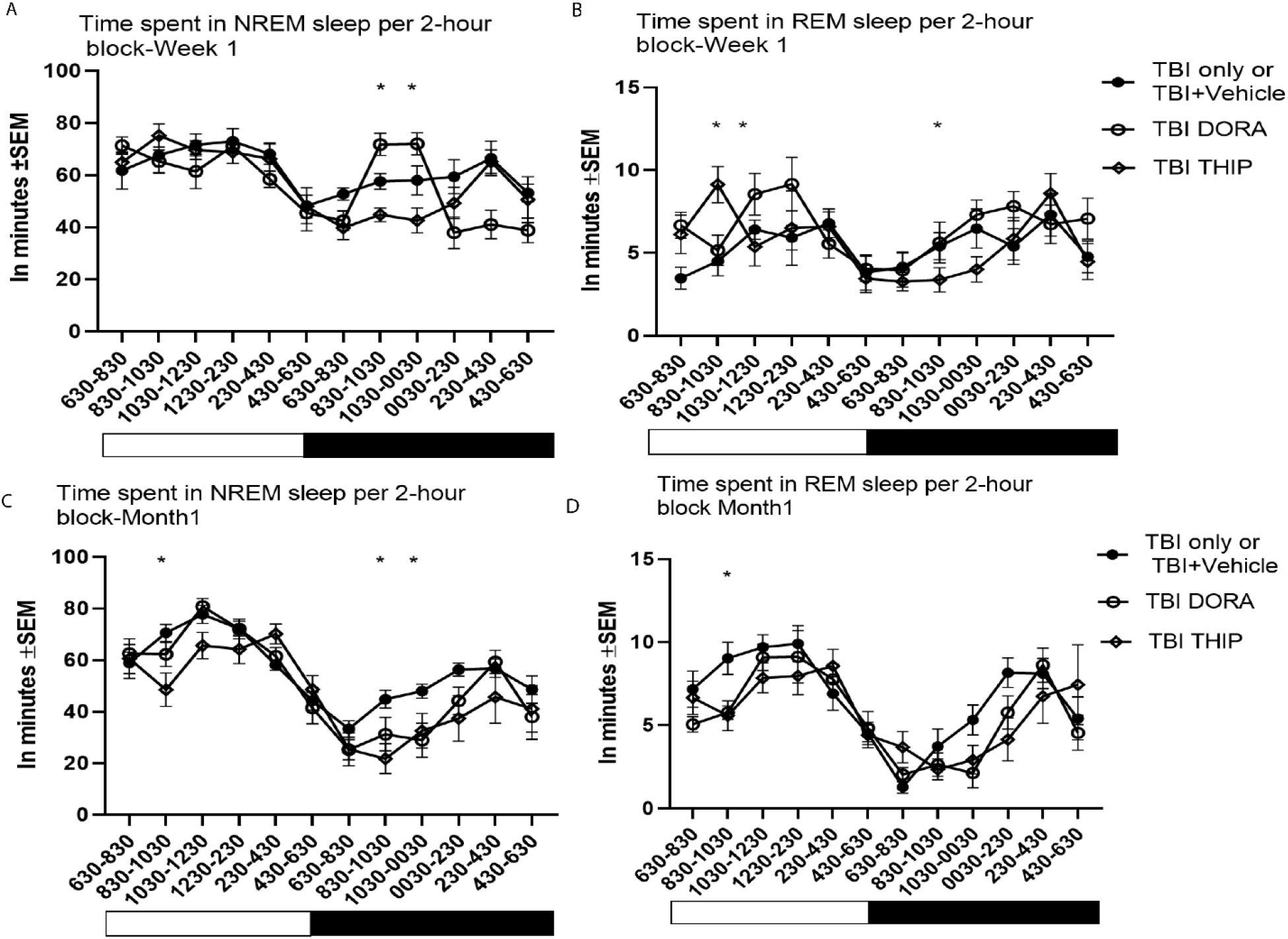
Sleep-wake analysis: The data for time spent in NREM at week 1 (A), month 1 (C), and REM sleep at week 1 (B) and month 1 (D) in each 2-hour block across lights on (light bar) and lights off (black bar) are shown along with the time indicated at the bottom. The normal diurnal oscillation of time in NREM or REM sleep is seen at both 1-week and 1-month recording time points. 2-way ANOVA showed an interaction of treatment and time for NREM at week 1 [F (22, 308) = 4.017; p<0.001] and month-1 [F (22, 264) = 1.848; p=0.01]. For REM sleep there was a significant interaction between treatment and time both at week 1 [F (22, 297) = 2.121; p=0.002) and month 1 [F (22, 286) = 1.582; p=0.04). Post hoc Tukey tests for multiple comparisons showed differences between groups at some of the recording time points where in week-1 DORA-22 increased time spent in NREM sleep during the first 4-hours of lights-off and REM sleep increased in first 4-hours of lights-on with both DORA-22 and THIP (indicated by *). At month-1 THIP treatment had lower time in NREM in the second 2-hour block of lights on and both treatments had lower time in NREM in two 2-hour block of lights-off. Both treatment groups had lower time in REM compared to TBI alone or vehicle in the second 2-hour block of lights-on (indicated by *).

### 4. TBI-induced increased sleep homeostatic drive was rescued by DORA-22 and THIP

Sleep NΔ is a marker of sleep homeostatic drive and the overnight decline in NΔ signifies the normal homeostatic process of decline in sleep pressure from the beginning of the night to the end of night^29^. We examined if NΔ, and its overnight decline is altered by TBI and its treatment with DORA-22 or THIP. We derived NΔ from frontal EEG from subsets of animals that had TBI alone or TBI with vehicle (n=12), and treated with DORA (n=8) OR THIP (n=8) as well as no craniotomy controls (n=4).

First, the mean NΔ for the entire 24-hour period was examined between different treatment groups. At week-1, all TBI groups inclusive of drug treated ones had higher mean 24-hour NΔ compared to no craniotomy control whereas sham groups were no different from no craniotomy controls (Figure 3A). At month-1 however, the mean 2-hour NΔ of TBI only with or without vehicle treatment remained higher than no craniotomy controls, whereas TBI groups treated with DORA or THIP was no different from no craniotomy controls (Figure 2B). We then examined the NΔ across the 24 hours in 1-hour blocks. Sham injury groups treated with DORA or THIP, at week 1 had no differences with each other (Figure 3 C) though at month 1 DORA treated sham group had higher NΔ that no craniotomy control at multiple time points (Figure 3 D). When TBI groups were examined separately, at week 1, all TBI groups (treated and untreated) had higher NΔ across every hour compared to no craniotomy whereas at month 1, no difference was seen between TBI groups treated with DORA and THIP, and no craniotomy control suggesting that the sleep aids restored the NΔ after TBI (Figure 3 E and F).

**Figure 3:**
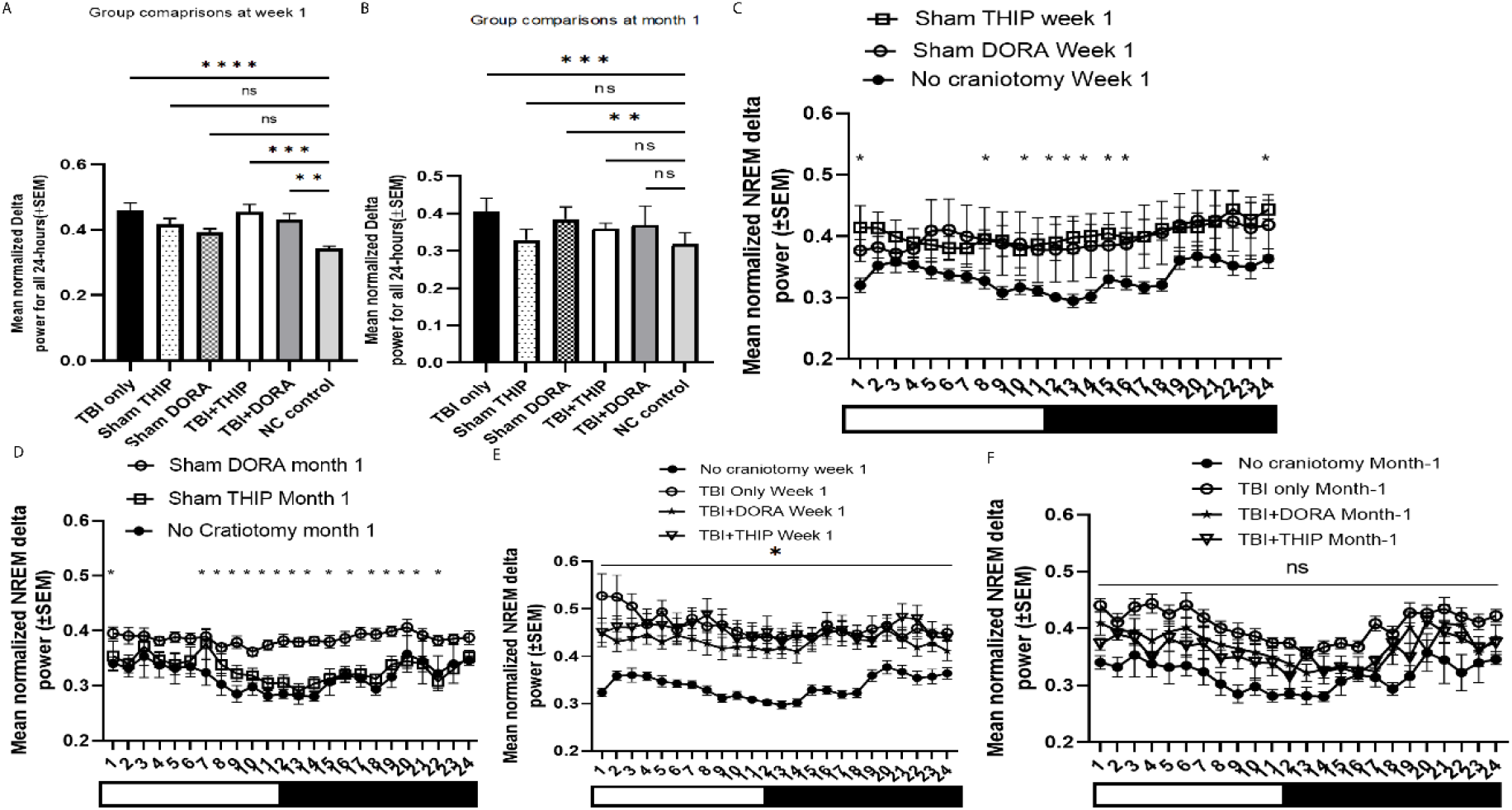
Data for NREM delta power at week 1 and month 1 is shown: <u>A and B</u>: The mean normalized delta power for the whole 24 hours of recording (±SEM) at week 1 was much higher in all TBI groups inclusive of drug treated ones compared to NC control [F (5, 56) = 7.578; p<0.001] (A) and at month-1 however, the mean normalized delta power of TBI groups treated with DORA or THIP was no different from NC control while the TBI only with or without vehicle treatment remained elevated [F (5, 45) = 6.497; p=0.001; post hoc Dunnett’s test showing only significant difference between TBI only and NC control; Figure 2B). <u>C and D</u>: The normalized delta power across the 24 hours in 1-hour blocks (±SEM) shown for sham injury groups treated with DORA or THIP at week 1 and month 1. At week 1, 2 way ANOVA showed no significant interaction between treatment group and time point [F (46, 621) = 0.9168; p=0.63; Figure 2C] though on post hoc Tukey tests for multiple comparisons, there were time points where drug treated sham animals had higher normalized delta power than no craniotomy controls. At month 1 there was a significant interaction of treatment and time point [F (46, 415) = 1.662; p=0.005; Figure 2D] with post hoc multiple comparisons showing that DORA treated sham group had higher NREM delta power at multiple time points (indicated by *). <u>E and F</u>: The normalized delta power (±SEM) showed for TBI groups that are untreated vs those treated with DORA and THIP at week 1 and month 1. At week 1, 2-way ANOVA showed a significant interaction of treatment and time point [F (69, 1012) = 2.292; p<0.001] and post hoc multiple comparisons showed that no craniotomy control has significantly lower NREM delta power at multiple time points (indicated by *). At month 1, no interaction between treatment group and time points was seen [F (69, 621) = 0.8372; p=0.82] and multiple comparisons showed no difference between TBI treated with DORA and THIP to that of no craniotomy control.

Finally, we examined whether the diurnal oscillation and overnight decline of NΔ is preserved in each of the treatment groups, by comparing NΔ in first 3 hours of lights-on (beginning of sleep) with the last 2 hours of lights-on (end of sleep and lights-off). No craniotomy controls had the normal homeostatic decline where it was higher in first 3 hours compared to last 2 hours of lights-on at week-1 and month-1 (Figure 4 A and E). TBI only or TBI plus vehicle, DORA and THIP treated CCI had loss of this homeostatic decline in week-1 (Figure 4 B-D). At month-1 the homeostatic decline or the oscillation of NΔ was restored in TBI only or TBI plus vehicle and DORA-22 treated groups but in the THIP treated TBI group this homeostatic decline of NΔ was lost (Figure 3 F-H).

**Figure 4:**
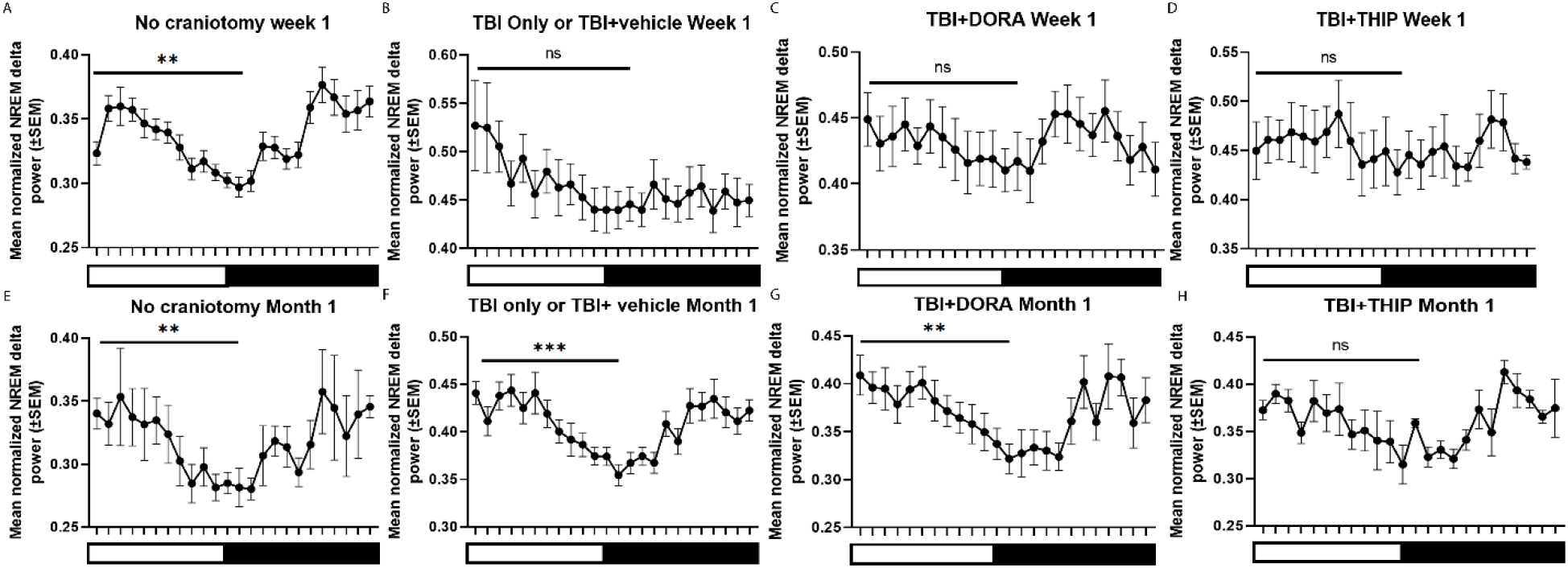
Figure shows the overnight decline in NΔ in week 1(A-D) and in month 1 (E-H). The mean normalized delta power was compared from the first part of the night (First 3 hours of lights-on) to the end of night (hours 11-12 of lights-on) between all the different treatment groups using paired t-tests and differences were indicated with asterisks when present (*= p<0.05; **= p<0.01; ***=p<0.001; ns=not significant).

## Discussion

The major and novel findings of our study are: a) Posttraumatic seizures are suppressed after the first week by a month long treatment with a sleep aid DORA (DORA-22) whereas similar treatment with another sleep aid THIP did not suppress seizures; b) TBI impaired GABAergic inhibition in dentate granule cells and is rescued by DORA-22 treatment; c) Sleep homeostatic drive as measured by NΔ increased markedly in the first week after TBI and remained high at the end of 1 month and treatment with both DORA-22 and THIP lowered NΔ to levels similar to animals that had no craniotomy and no TBI; and d) TBI resulted in loss of the normal homeostatic overnight decline in NΔ but this was restored at 1 month post-TBI in the TBI only or TBI with vehicle and DORA-22 treatment groups but not in the THIP treatment group.

Our data on seizure suppressing effects of DORA treatment is consistent with prior reports in mouse models of epilepsy^19-21^. The exact mechanisms of the anti-seizure effects of orexin antagonists had not been studied before. However, orexin-A injections into hippocampus or into CSF was shown to increase glutamate release whereas orexin antagonists reduced glutamate^30^. In line with this, orexin antagonists decreased glutamate levels and increased GABA levels in the hippocampus after pentylenetetrazol induced seizures^31^. Our data showed that TBI decreased frequency and amplitude of mIPSCs in dentate granule cells and DORA-22 increased them suggesting that DORA-22 treatment enhances GABAergic inhibition and may be a potential mechanism by which DORA-22 treatment suppressed posttraumatic seizures.

EEG delta power in NREM sleep has been used as a proxy or marker for sleep homeostatic drive, wherein the slow wave activity or delta power in NREM sleep increases proportionately with increasing wake duration and declines progressively with subsequent sleep^23^, which was established from sleep deprivation studies^32^. According to the synaptic homeostasis hypothesis, the normal overnight decline of sleep slow wave or delta activity in NREM sleep across the night serves to prune or weaken synapses strengthened during prior wake which is important for learning and memory^33^. Prior studies indicated that the increased slow wave activity or delta power during sleep is triggered by an increase in synaptic strength (i.e., synaptic potentiation)^34^. The overnight decrease in delta activity in NREM sleep and NΔ is thought to reflect the net effect of synaptic down-selection occurring during NREM sleep^35^ which makes synapses more available for new learning during awake state that follows sleep. In an epileptic brain, a prior study showed that the NΔ is elevated both globally in the brain and locally in the seizure onset zones and a lack of normal overnight decline or the loss of normal oscillation across the night correlated with memory impairments^36^. In the mouse model of TBI, We found that the 24-hour mean NΔ increases both acutely similar to other reports^37^ but also chronically^13^ and it is possible that TBI induces synaptic potentiation similar to that of an epileptic brain^38^. Moreover, the mean NΔ that increased after TBI (no drug or vehicle treatment) remained elevated at 1-month suggesting that synaptic potentiation can be long lasting. Interestingly we found that a month-long treatment with sleep aids DORA-22 and THIP lowered the mean NΔ across the 24-hours bringing it to levels of animals that had no craniotomy. Our data are consistent with another study which showed that treatment with DORA-22 lowered NΔ significantly at a dose of 30mg/kg in a mouse model of insomnia but also improved memory^39^. It is possible that the reduction of the NΔ is favorable for synaptic plasticity and memory function^40^. While THIP lowered NΔ as well, it did not prevent PTE and prior studies showed that THIP does not necessarily suppress seizures^41^. THIP and DORA were shown to increase NΔ in normal animals^42-43^ but we found a reduction after month-long treatment. Whether reducing NΔ following TBI or in an epileptic brain drives seizure suppression remains to be explored further.

We also found that the normal homeostatic decline in NΔ is disrupted at least acutely after TBI in the first week which had not been previously reported. The loss of this homeostatic decline is unfavorable for normal synaptic plasticity and can contribute to memory problems. Treatment with THIP failed to restore this normal decline in NΔ even chronically suggesting that this drug may be associated with impairments in learning and memory as reported previously^44^. DORA-22 on the other hand restored it and may be favorable from a memory standpoint during the subsequent day^39^.

In normal animals, DORAs are kwon to increase time spent in NREM and REM sleep^45^. In another study, DORA-22 on the other hand increased time spent in NREM sleep in the first 3 hours but not in the second 3-hours^38^. Interestingly, the time spent in sleep or awake across the entire 24-hours did not increase with DORA-22 in a mouse model of Alzheimer’s Disease^46^ or insomnia^39^. Our data are consistent with these latter reports and we did not find an increase in time spent in NREM or REM sleep when the entire 24-hours or 12-hour blocks of lights-on and lights-off were considered.

There are several limitations to highlight in our study. First, we did not perform continuous EEG recordings for all 3-months of the study period and it is possible that we missed seizures during the time we did not record. Future studies should perform continuous EEG recordings for the duration of the experiment to better understand effects of DORAs on seizures and sleep after TBI^18^. We did not analyze spindles and DORA-22 is shown to increase spindle number in the first hour of post-treatment sleep^39^. We have not analyzed interictal spikes which may have influence on NΔ. We have limited electrophysiology data only from dentate granule cells and future studies should perform a more comprehensive study examining both excitatory and inhibitory neurotransmission in the hippocampus, cortex and possibly the thalamus which is involved in generation of NREM delta activity. Finally, future studies should also employ histopathological examination to more conclusively understand whether DORA-22 can have disease modifying effect for PTE.

## Competing Interests and Disclosures

None to declare

## Funding Source

Department of Defense Grant PRMRP:161864 (PI: RM) and National Institutes of Health Grant Number R21NS104612-01A1 (PI:RM) and R21NS116546 (PI:MVJ).

## Notes

### Competing Interest Statement

The authors have declared no competing interest.

